# Effects of sex, mating status, and genetic background on circadian behavior in *Drosophila*

**DOI:** 10.1101/2024.11.22.624853

**Authors:** Oghenerukevwe Akpoghiran, Alexandra K. Strich, Kyunghee Koh

## Abstract

Circadian rhythms play a crucial role in regulating behavior, physiology, and health. Sexual dimorphism, a widespread phenomenon across species, influences circadian behaviors. Additionally, post-mating physiological changes in females are known to modulate various behaviors, yet their effects on circadian rhythms remain underexplored. Here, using *Drosophila melanogaster*, a powerful model for studying circadian mechanisms, we systematically assessed the impact of sex and mating status on circadian behavior. We measured circadian period length and rhythm strength in virgin and mated males and females, including females mated to males lacking Sex Peptide (SP), a key mediator of post-mating changes. Across four wild-type and control strains, we found that males consistently exhibited shorter circadian periods than females, regardless of mating status, suggesting that circadian period length is a robust sexually dimorphic trait. In contrast, rhythm strength was influenced by the interaction between sex and mating status, with female mating generally reducing rhythm strength in the presence of SP signaling. Notably, genetic background significantly modulated these effects on rhythm strength. Our findings demonstrate that while circadian period length is a stable sex-specific trait, rhythm strength is shaped by a complex interplay between sex, mating status, and genetic background. This study advances our understanding of how sex and mating influence circadian rhythms in *Drosophila* and provides a foundation for future research into sexually dimorphic mechanisms underlying human diseases associated with circadian disruptions.

## 1 Introduction

Circadian rhythms are approximately 24-hour cycles in behavior and physiology generated by endogenous molecular clocks that are highly conserved across species (Creux and Harmer, 2019; Dunlap and Loros, 2017; Hardin, 2011; Johnson et al., 2017; Lane et al., 2023). These clocks function through transcriptional-translational feedback loops, where activators repress their own activation over a 24-hour cycle (Allada and Chung, 2010; Patke et al., 2020). The circadian clock integrates environmental inputs to regulate various rhythmic processes. Disruptions in these rhythms are linked to sleep, metabolic, and neurodegenerative disorders, highlighting the importance of studying circadian rhythms (Leng et al., 2019; Sack et al., 2007; Shimizu et al., 2016).

Sexual dimorphism is widespread, with males and females differing in morphology, physiology, and behavior (Williams and Carroll, 2009). Sex differences in circadian patterns are also common across species (Joye and Evans, 2022). For example, women generally go to bed earlier than men, and pregnancy shortens women’s circadian period (Martin-Fairey et al., 2019; Roenneberg et al., 2007). Given the conservation of the circadian clock across species, *Drosophila melanogaster* is an excellent model organism for studying circadian rhythms. Previous research has shown that male and female flies exhibit distinct circadian behaviors (Helfrich-Förster, 2000). Males show more pronounced evening activity, while females display higher overall activity levels throughout the day. Additionally, male flies have shorter circadian periods than female flies. Most studies of *Drosophila* circadian behavior employ males partly due to anecdotal observations suggesting that males have more robust rhythms than females; however, systematic investigations into sex differences in rhythm strength are lacking.

Beyond sex differences, mating induces notable behavioral and physiological changes in female *Drosophila*. During copulation, males transfer Sex Peptide (SP) to females (Chen et al., 1988; Liu and Kubli, 2003), triggering various behavioral changes, including increased egg laying, heightened food intake, and sleep loss (Kubli and Bopp, 2012). Additionally, a recent study found that female mating disrupts morning anticipation, which refers to the anticipatory activity before dawn, under light:dark (LD) conditions through SP signaling (Riva et al., 2022). However, a prior study found no significant effect of female mating on circadian period length in constant darkness (DD) (Helfrich-Förster, 2000), and the impacts of female mating status and SP on circadian rhythm strength remain unexplored, as does the effect of male mating status on circadian behavior.

To investigate the roles of sex, mating status, and genetic background in *Drosophila* circadian behavior, we studied four wild-type and control strains, assessing circadian period length and rhythm strength in virgin males, mated males, virgin females, females mated with same-strain males, and females mated with SP-deficient males. This approach allowed us to explore the effects of sex, mating status, SP, and genetic background on circadian locomotor activity. Our data show that circadian period regulation is sexually dimorphic across strains, with males exhibiting shorter circadian periods than females regardless of mating status and genetic background. In contrast, our findings indicate that the effects of sex and mating status on rhythm strength are strongly modulated by genetic background.

## 2 Materials and methods

### Fly Stocks

Flies were raised on standard food containing cornmeal, molasses, and yeast at 25°C and under a 12h:12h light-dark (LD) cycle. The following lines were obtained from the Bloomington *Drosophila* Stock Center: *Canton-S* (#64349), *iso31* (*w*^1118^, #3605), *Oregon-R-modENCODE* (#25211), *Oregon-R-P2* (#2376), and *SP*^0^ (#94694).

### Circadian Assays

Virgin males and females were collected shortly after eclosion and kept in groups of ∼32 for 2 to 4 days in a 25°C LD cycle. Approximately 1 day before the circadian assays, 16 virgin females were placed with 16 virgin males of the same strain in each vial to produce mated males and females mated within strain. Additionally, virgin females of all strains were crossed to *SP*^0^ mutant males. Flies in the virgin condition were transferred to fresh vials when the flies in the mated condition were crossed.

We utilized the *Drosophila* Activity Monitoring (DAM) System (TriKinetics) to record circadian locomotor activity over approximately 6.5 days in DD. Flies were placed individually in small glass tubes containing 5% sucrose and 2% agar. Data spanning 6 days in DD were analyzed to calculate rhythm parameters. We determined the circadian period length between 18 and 30 h and rhythm strength for each fly using Faas software (https://neuropsi.cnrs.fr/en/departments/cnn/group-leader-francois-rouyer/). This software utilizes χ2 periodogram analysis on activity counts recorded in 30-minute intervals. Rhythm strength is defined as the difference between the peak χ2 value and the expected χ2 value at chance (p = 0.05). Flies were categorized by rhythm strength into three groups: rhythmic (>50), weakly rhythmic (25–50), and arrhythmic (25). Arrhythmic flies were excluded from the analysis of period length because accurately determining a circadian period requires a sufficiently robust rhythm strength. For rhythm strength analysis, however, all flies were included to provide a comprehensive view of rhythmicity within each group. This approach ensured a single, unified metric of rhythmicity that accounts for both the proportion of rhythmic individuals and the average rhythm strength of rhythmic flies, simplifying the assessment.

## Statistical Analysis

Prism 10 (GraphPad Software) was used for statistical analysis. Multiple group comparisons were performed using Brown-Forsythe and Welch’s ANOVA tests followed by Dunnett’s T3 multiple comparisons tests.

## 3 Results

### Males exhibit shorter period lengths than females

We employed the *Drosophila* activity monitor system to examine the free-running locomotor behavior of virgin and mated flies of both sexes in DD. Four strains were included: *Canton S* (a widely used wild-type strain), *iso31* (an isogenic strain carrying the *w*^1118^ mutation in the *Canton-S* background), *Oregon-R modENCODE* (an *Oregon-R* variant utilized by the modENCODE project (The modENCODE Consortium et al., 2010)), and *Oregon-R-P2* (an *Oregon-R* strain known for rapid egg-laying (Mahowald et al., 1983)). Female flies were either mated with males of their own strain or SP-deficient *SP*^0^ mutant males. Notably, males in all strains exhibited significantly shorter circadian periods than females, with differences of ∼0.2 h in iso31 and ∼0.4–0.6 h in the other strains (Fig. 1 and Table 1). This consistent sexual dimorphism was observed across all strains despite variations in mean period length between strains; the mean period in males was ∼24.5 h for Canton-S, ∼23.5 h for iso31, and ∼23 h for both Oregon-R-modENCODE and Oregon-R-P2, with females showing a similar trend.

**Table 1:** Circadian locomotor rhythm phenotypes of four *Drosophila* strains in DD.

| **Genotype** | **N** | **%R** | **%WR** | **%AR** | **tau ± SEM** | **Power ± SEM** |
| --- | --- | --- | --- | --- | --- | --- |
| ***Canton-S*** |  |  |  |  |  |  |
| Virgin males | 81 | 97.5 | 3.7 | 2.5 | 24.54 ± 0.05 | 119 ± 3.48 |
| Mated males | 104 | 95.2 | 3.8 | 4.8 | 24.49 ± 0.03 | 110.8 ± 3.63 |
| Virgin females | 99 | 98.0 | 5.1 | 2.0 | 24.96 ± 0.04 | 109.8 ± 3.29 |
| SP<sup>0</sup> mated females | 59 | 96.6 | 1.7 | 3.4 | 25.07 ± 0.06 | 107.7 ± 4.86 |
| Mated females | 92 | 96.7 | 9.8 | 3.3 | 24.89 ± 0.04 | 88.06 ± 3.25 |
| ***iso31*** |  |  |  |  |  |  |
| Virgin males | 102 | 93.1 | 5.9 | 6.9 | 23.54 ± 0.02 | 112.3 ± 4.46 |
| Mated males | 134 | 96.3 | 1.5 | 3.7 | 23.57 ± 0.02 | 115.3 ± 3.50 |
| Virgin females | 123 | 98.4 | 2.4 | 1.6 | 23.73 ± 0.04 | 120.7 ± 3.32 |
| SP<sup>0</sup> mated females | 104 | 99.0 | 5.8 | 1.0 | 23.75 ± 0.04 | 114.9 ± 3.66 |
| Mated females | 130 | 93.1 | 15.4 | 6.9 | 23.79 ± 0.03 | 81.02 ± 3.45 |
| ***Oregon-R-modENCODE*** |  |  |  |  |  |  |
| Virgin males | 165 | 78.8 | 26.7 | 21.2 | 22.97 ± 0.04 | 57.01 ± 2.86 |
| Mated males | 173 | 74.6 | 20.2 | 25.4 | 22.93 ± 0.06 | 53.12 ± 2.85 |
| Virgin females | 153 | 95.4 | 5.9 | 4.6 | 23.43 ± 0.03 | 110.5 ± 3.78 |
| SP<sup>0</sup> mated females | 147 | 98.0 | 16.8 | 2.0 | 23.46 ± 0.03 | 107.1 ± 3.61 |
| Mated females | 167 | 94.0 | 6.8 | 6.0 | 23.46 ± 0.02 | 80.36 ± 2.86 |
| ***Oregon-R-P2*** |  |  |  |  |  |  |
| Virgin males | 117 | 98.3 | 3.4 | 1.7 | 23.04 ± 0.03 | 131.9 ± 3.82 |
| Mated males | 158 | 96.2 | 2.5 | 3.8 | 23.21 ± 0.05 | 122.3 ± 3.43 |
| Virgin females | 137 | 97.1 | 3.6 | 2.9 | 23.66 ± 0.03 | 134.7 ± 3.54 |
| SP<sup>0</sup> mated females | 139 | 99.3 | 2.2 | 0.7 | 23.72 ± 0.04 | 134.9 ± 3.14 |
| Mated females | 147 | 98.0 | 1.4 | 2.0 | 23.99 ± 0.05 | 131.0 ± 2.85 |

N, number of flies; R, rhythmic; WR, weakly rhythmic; AR, arrhythmic. tau, free-running period; power, a measure of rhythm strength.

**Figure 1:**
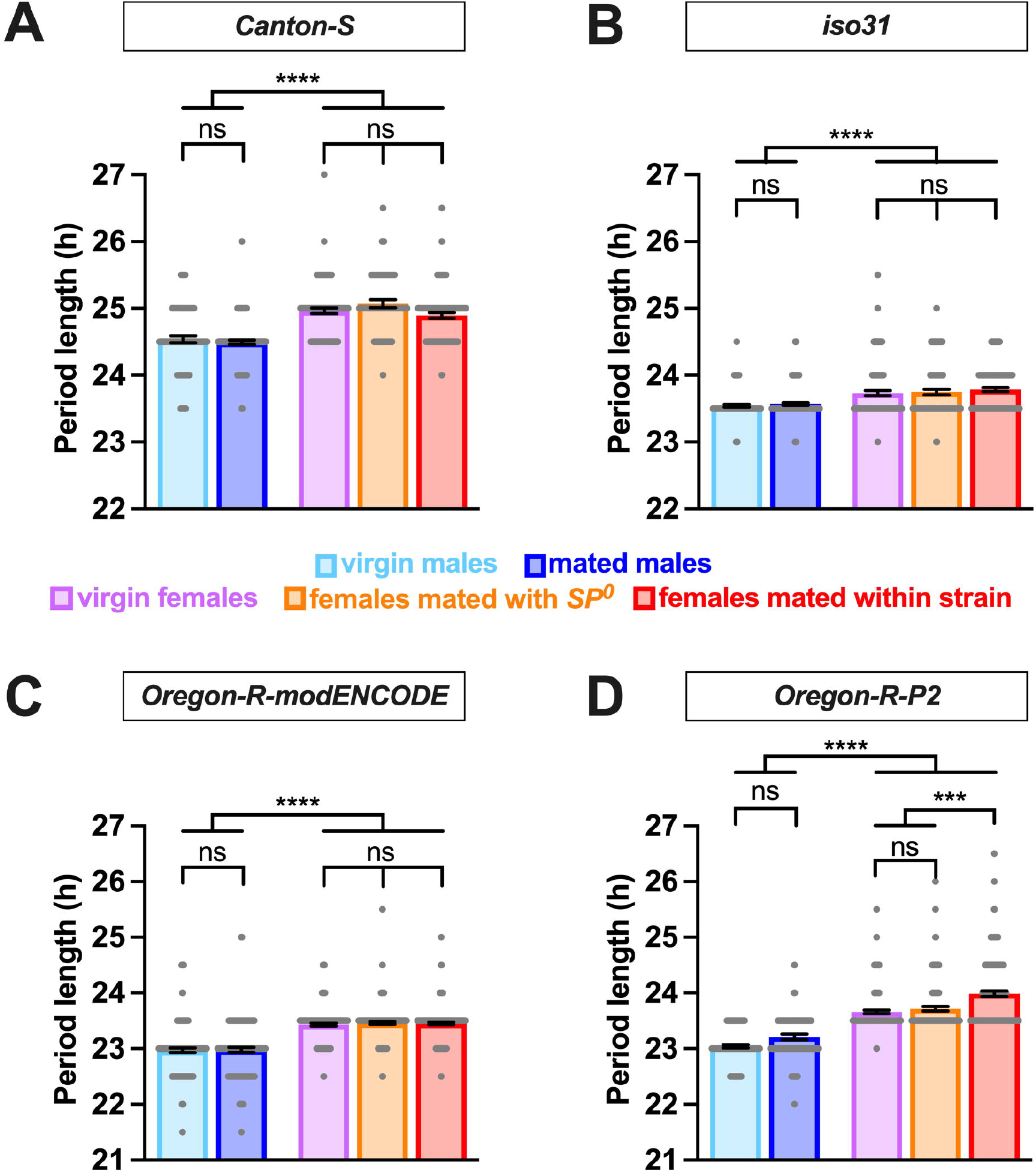
Male flies have a shorter circadian period length than females. The circadian period length of *Canton-S* (A), *iso31* (B), *Oregon-R-modENCODE* (C), and *Oregon-R-P2* (D) is shown. The mean and SEM are shown. N = 57 -157. ***p < 0.001, ****p < 0.0001, ns, not significant; Brown-Forsythe and Welch’s ANOVA tests followed by Dunnett’s T3 multiple comparisons.

Whereas sex had a significant and consistent effect on the circadian period length, mating status had minimal impact. In all strains, there was no significant difference in period length between virgin and mated males (Fig. 1 and Table 1). Similarly, females of different mating statuses showed comparable circadian periods, with a minor exception in the *Oregon-R-P2* strain, where females mated within strain had significantly longer periods than virgin and *SP*^0^-mated females. However, in all strains, females consistently exhibited longer periods than males, regardless of the mating status.

Overall, these findings suggest that circadian period regulation in *Drosophila* is influenced by sex but is minimally impacted by mating status and genetic background.

### Effects of sex and mating on rhythm strength are modulated by genetic background

We also examined how sex, mating status, SP, and genetic background affect the strength of circadian rhythmicity. While circadian period length showed consistent sex differences across strains, the effects of sex and mating status on rhythm strength was more variable across strains. In *Canton S* and *iso31*, rhythm strength was similar among virgin males, mated males, virgin females, and *SP*^0^-mated females (Fig. 2A-B and Table 1), indicating that sex alone does not influence rhythm strength in these strains. However, females mated within strain showed a marked reduction in rhythm strength compared to virgin and *SP*^0^-mated females, suggesting that female mating status modulates their rhythm strength via the SP signaling pathway.

**Figure 2:**
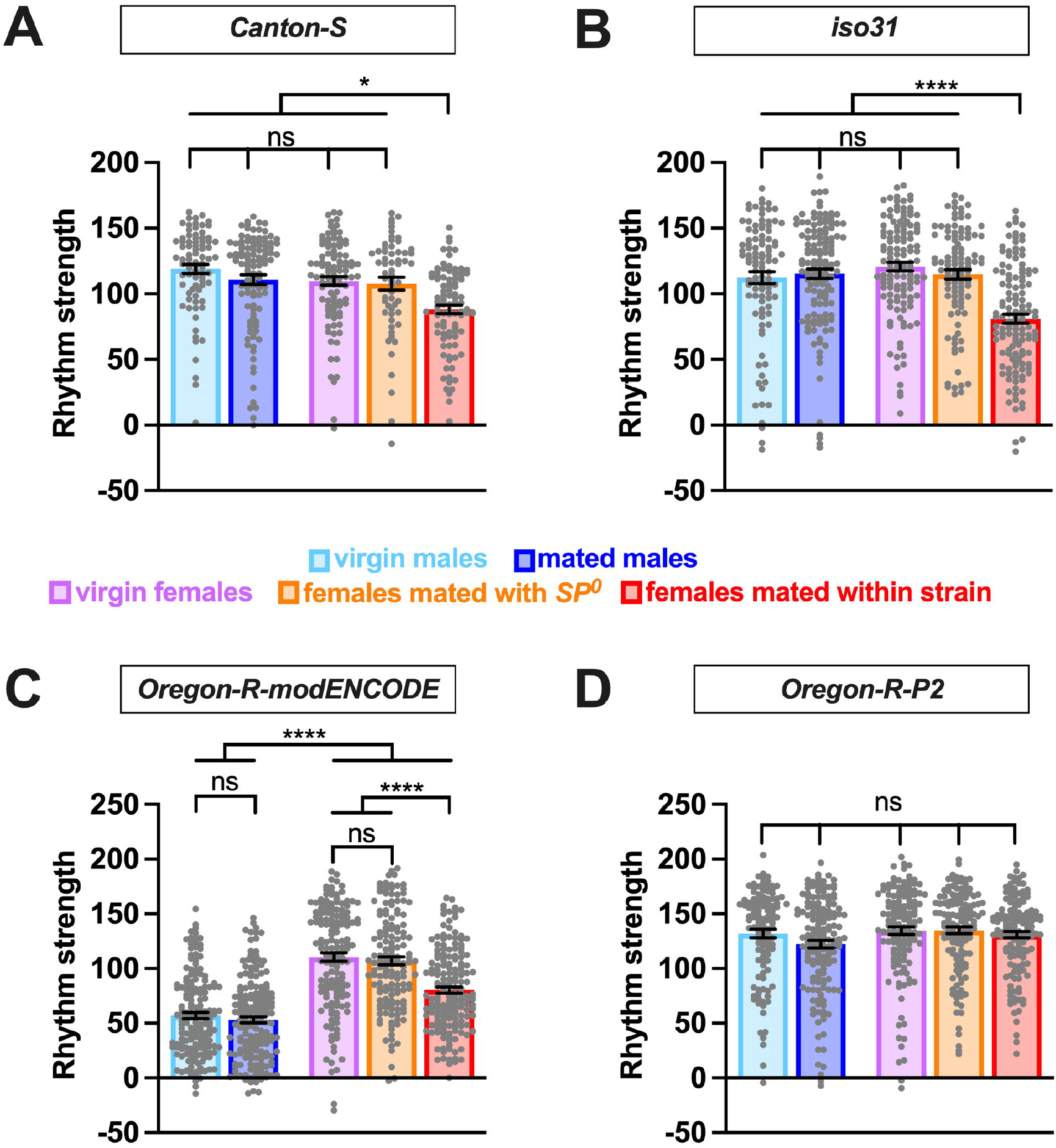
Effects of sex and mating on circadian rhythm strength are modulated by genetic background. The rhythm strength of *Canton-S* (A), *iso31* (B), *Oregon-R-modENCODE* (C), and *Oregon-R-P2* (D) is shown. The mean and SEM are shown. N = 59 -173. *p < 0.05; ****p < 0.0001; ns, not significant; Brown-Forsythe and Welch’s ANOVA tests followed by Dunnett’s T3 multiple comparisons.

Similar to *Canton S* and *iso31, Oregon-R-modENCODE* females mated within strain exhibited significantly lower rhythm strength than virgin and *SP*^0^-mated females (Fig. 2C and Table 1). Unexpectedly, males in this strain displayed significantly lower rhythm strength than females, with 21-26% arrhythmic males versus only 2-6% arrhythmic females (Table 1). This suggests that both sex and the female mating status influence rhythm strength in *Oregon-R-modENCODE*. In contrast, for *Oregon-R-P2*, neither sex nor mating status had a significant effect on rhythm strength (Fig. 2D and Table 1), as all pairwise comparisons were non-significant. Unlike the other strains, female mating status did not affect rhythm strength in *Oregon-R-P2*.

Together, these findings suggest that females generally exhibit reduced rhythm strength after mating with wild-type SP-containing males. Additionally, genetic background has a stronger influence on rhythm strength than on circadian period length.

## 4 Discussion

We investigated how sex, mating status, and genetic background influence the period length and strength of free-running rhythms in DD in *Drosophila melanogaster*. Our data show that these factors impact circadian period length and rhythm strength in distinct ways. Males consistently had shorter circadian periods than females, regardless of mating status or genetic background, revealing a robust sexual dimorphism in the speed of the *Drosophila* circadian clock. However, rhythm strength was modulated by a complex interaction among sex, mating status, and genetic background. In three strains (*Canton-S, iso31*, and *Oregon-R-modENCODE*), females mated with *SP*^*+*^ males showed reduced rhythm strength compared to virgin females and *SP*^0^-mated females, with *Oregon-R-P2* being the exception. Except for *Oregon-R-modENCODE*, males exhibited rhythm strength similar to virgin and *SP*^0^-mated females. These findings suggest that rhythm strength is primarily influenced by female mating status and genetic background rather than inherent sexual dimorphism. While sexual dimorphism in circadian period length is independent of mating status and genetic background, the effects of sex on rhythm strength are modulated by mating status and genetic background. This difference may arise because the circadian period is determined by the pace of the molecular clock in clock neurons, while rhythm strength is influenced by additional output mechanisms in both clock and downstream neurons, making it more susceptible to modulation by genetic background and female mating status.

Several studies have identified sexually dimorphic gene expression and function in clock neurons in *Drosophila*. The sex-specific neural circuitry and social behaviors in *Drosophila* are orchestrated by the interplay of sexually dimorphic transcription factors encoded by *fruitless (fru)* and *doublesex (dsx*) (Ellendersen and von Philipsborn, 2017; Sato and Yamamoto, 2023; Yamamoto and Koganezawa, 2013). Previous research has shown that certain clock neuron clusters contain cells expressing a male-specific isoform of *fru* (Fujii and Amrein, 2010), and that a greater number of clock neurons in a dorsal neuron 1 (DN1) cluster express *fru* in males compared to females (Hanafusa et al., 2013). A recent transcriptomic analysis of *fru*-expressing neurons using single-cell RNA sequencing identified a specific set of *fru-*expressing clock neurons (Palmateer et al., 2023). Interestingly, artificial activation of these neurons had sexually dimorphic effects on the circadian period; males exhibited a shortened period, while females displayed a lengthened period. These findings suggest that *fru*-expressing clock neurons contribute to the sex differences observed in the present study. Additionally, a subset of dorsal lateral neurons (LNds) shows male-specific expression of Neuropeptide F (NPF) (Hermann et al., 2012; Lee et al., 2006), which is associated with feeding, metabolism, and courtship (Cui and Zhao, 2020). Another neuropeptide, Pigment Dispensing Factor (PDF) is crucial for synchronizing rhythms in DD (Renn et al., 1999), and the absence of PDF or its receptor, PDFR, leads to arrhythmic activity-rest cycles in most male flies (Lear et al., 2005; Mertens et al., 2005; Peng et al., 2003; Yoshii et al., 2009). Intriguingly, a recent study found that mutations in *Pdf* and *Pdfr*, have a lesser impact on female circadian rhythms than on males, suggesting a sexually dimorphic role for PDF in circadian regulation (Iyer et al., 2024). The molecular mechanisms by which sex differences in neural circuits and gene expression translate into distinct circadian behaviors in *Drosophila* warrant further investigation.

Previous studies have documented several post-mating changes in *Drosophila* females, including increased egg-laying, decreased sexual receptivity, and reduced sleep (Kubli and Bopp, 2012). These changes are primarily driven by SP transferred from males during copulation (Chen et al., 1988; Liu and Kubli, 2003). Our data add reduced rhythm strength to the list of SP-dependent post-mating behavioral changes. Interestingly, the post-mating reduction in rhythm strength was absent in the *Oregon-R-P2* strain, implying that unique genetic factors in this variant counteract the typical post-mating changes. *Oregon-R-P2* females are known for rapid egg-laying (Lynch et al., 1989), hinting that the post-mating reduction in rhythmicity observed in the other strains may be linked to normal egg-laying. Additionally, recent studies showed that mated females exhibit significantly lower PDF levels compared to virgin females (Riva et al., 2022), and mating and SP trigger changes in the expression of core clock genes along with a loss of rhythmicity in many clock-controlled genes (Delbare et al., 2023). These molecular changes may underlie the post-mating reduction in rhythm strength.

We recently reported that female post-mating reduction in nighttime sleep depends on diet rich in nutrients, including protein, and is absent on 5% sucrose food (Duhart et al., 2023). Diet may also modulate circadian behavior. However, it is challenging to examine the effects of mating on circadian behavior under protein-rich conditions due to the rapid development of larvae, which can disturb behavioral recordings within a few days. Consequently, in our experiments, flies were raised on a nutrient-rich diet but were housed in tubes containing 5% sucrose during the circadian assays. Interestingly, a recent study investigated the effects of sex and mating status on morning anticipation using a protein-rich medium under LD conditions (Riva et al., 2022). Morning anticipation in LD can be assessed within a few days of mating before larval movement becomes problematic. The study found that mated females exhibited reduced morning anticipation compared to virgin females and males, which may be linked to a decrease in rhythm power. How mating influences the circadian period and rhythm strength in DD under nutrient-rich conditions is an interesting but technically challenging topic for future investigation.

Sex differences in circadian behaviors are common across species (Bertossa et al., 2013; Helfrich-Förster, 2000; Walton et al., 2022). For example, in rodents, some studies showed that females generally have shorter endogenous period lengths than males, although this sex difference did not always reach statistical significance (Schull et al., 1989; Sterniczuk et al., 2010;Iwahana et al., 2008; Kuljis et al., 2013). Similarly, women tend to go to bed earlier than men from childhood to menopause, at which time sex differences in sleep patterns disappear (Roenneberg et al., 2007).

Moreover, a recent study reported that pregnant women had phase-advanced endogenous temperature and melatonin rhythms (Cain et al., 2010), partly because of a significantly shorter circadian period length (Duffy et al., 2011). While the effects of sex and pregnancy on circadian periods in humans and rodents are well documented, their impact on circadian rhythm strength remains an open question.

Interestingly, whereas males exhibit longer periods than females in humans and rodents, female flies display longer periods than male flies. Species-specific behaviors may underlie these distinct sexual dimorphisms across organisms. For example, a shorter circadian period in male flies may allow them to wake up earlier, making it easier for them to find and court inactive females rather than active ones (Helfrich-Förster, 2000). Investigating how sexually dimorphic tasks influence the molecular clock and circadian behavior may advance our understanding of sex-specific regulation of biological rhythms.

Neurodegenerative disorders, including Parkinson’s disease, Alzheimer’s disease, Amyotrophic Lateral Sclerosis (ALS), cardiovascular disease, and sleep disorders, are associated with circadian disruptions, which may exacerbate these disease symptoms (Fifel and Videnovic, 2021; Videnovic et al., 2014). Additionally, these diseases display sexual dimorphism (Curtis et al., 2017; Haaxma et al., 2007; McCombe and Henderson, 2010; Oveisgharan et al., 2018). Unraveling the genetic and neural factors that drive sex-specific variations in circadian rhythms may provide a broader understanding of circadian misalignment in disease contexts and suggest targeted interventions for improving health outcomes.

## Author Contributions

Conceptualization & Methodology, O.A and K.K. Investigation, O.A and A.S. Writing – Original Draft, O.A and K.K. Writing – Review & Editing, A.S. Funding Acquisition & Supervision, K.K.

### Funding

This work was supported by grants from the National Institutes of Health (R01NS086887 and R01NS109151 to K.K.) and funds from the Thomas Jefferson University Synaptic Biology Center (to K.K.).

## Acknowledgments

We thank the Bloomington Drosophila Stock Center for fly stocks; M. Boudinot and Dr. Francis Rouyer for the Faas software; Drs. Yohei Kirino, Joanna Chiu, Aaron Haeusler, and members of the Koh lab for helpful discussion.

